# Microglia Morphological Response to Mesenchymal Stromal Cell Extracellular Vesicles Demonstrates EV Therapeutic Potential for Modulating Neuroinflammation

**DOI:** 10.1101/2024.07.01.601612

**Authors:** Kanupriya R. Daga, Andrew M. Larey, Maria G. Morfin, Kailin Chen, Sara Bitarafan, Jana M. Carpenter, Hannah M. Hynds, Kelly M. Hines, Levi B. Wood, Ross A. Marklein

## Abstract

**Background:** Mesenchymal stromal cell derived extracellular vesicles (MSC-EVs) are a promising therapeutic for neuroinflammation. MSC-EVs can interact with microglia, the resident immune cells of the brain, to exert their immunomodulatory effects. In response to inflammatory cues, such as cytokines, microglia undergo phenotypic changes indicative of their function e.g. morphology and secretion. However, these changes in response to MSC-EVs are not well understood. Additionally, no disease-relevant screening tools to assess MSC-EV bioactivity exist, which has further impeded clinical translation. Here, we developed a quantitative, high throughput morphological profiling approach to assess the response of microglia to neuroinflammation-relevant signals and whether this morphological response can be used to indicate the bioactivity of MSC-EVs.

**Results:** Using an immortalized human microglia cell-line, we observed increased size (perimeter, major axis length) and complexity (form factor) upon stimulation with interferon-gamma (IFN-γ) and tumor necrosis factor-alpha (TNF-α). Upon treatment with MSC-EVs, the overall morphological score (determined using principal component analysis) shifted towards the unstimulated morphology, indicating that MSC-EVs are bioactive and modulate microglia. The morphological effects of MSC-EVs in TNF-γ/IFN-α stimulated cells were concomitant with reduced secretion of 14 chemokines/cytokines (e.g. CXCL6, CXCL9) and increased secretion of 12 chemokines/cytokines (e.g. CXCL8, CXCL10). Proteomic analysis of cell lysates revealed significant increases in 192 proteins (e.g. HIBADH, MEAK7, LAMC1) and decreases in 257 proteins (e.g. PTEN, TOM1, MFF) with MSC-EV treatment. Of note, many of these proteins are involved in regulation of cell morphology and migration. Gene Set Variation Analysis revealed upregulation of pathways associated with immune response, such as regulation of cytokine production, immune cell infiltration (e.g. T cells, NK cells) and morphological changes (e.g. Semaphorin, RHO/Rac signaling). Additionally, changes in microglia mitochondrial morphology were measured suggesting that MSC-EV modulate mitochondrial metabolism.

**Conclusion:** This study comprehensively demonstrates the effects of MSC-EVs on human microglial morphology, cytokine secretion, cellular proteome, and mitochondrial content. Our high-throughput, rapid, low-cost morphological approach enables screening of MSC-EV batches and manufacturing conditions to enhance EV function and mitigate EV functional heterogeneity in a disease relevant manner. This approach is highly generalizable and can be further adapted and refined based on selection of the disease-relevant signal, target cell, and therapeutic product.

## 1. Background

Mesenchymal stromal cell derived-extracellular vesicles (MSC-EVs) have been investigated for their therapeutic potential across a wide range of diseases (1–3). Their function is mediated through delivery of cargo, such as proteins, lipids and miRNAs (4,5). Compared to the MSCs that produce them, MSC-EVs may be safer due to reduced risks of thrombosis and tumorigenicity (6–12). Additionally, EVs derived from other cell types have been shown to readily cross physiological barriers (e.g. blood-brain-barrier) in the body due to their small size and targeting capability (13–18). Multiple *in vitro* and *in vivo* studies have shown their safety and therapeutic efficacy in different neurodegenerative disease models (4,19).

Despite these advantages, there are several challenges with using MSC-EVs as therapeutics. Like MSCs, MSC-EVs have well-recognized functional heterogeneity, which is similarly derived from differences in cell source (donor and tissue) and a lack of standardized manufacturing conditions (20). The International Society of Extracellular Vesicles (ISEV) criteria for EV characterization, like size, particle count, protein content, presence of tetraspanins (CD63, CD81 and CD9), and single vesicle analysis, are limited to particle identification (21); however, they do not reflect the bioactivity (i.e. functional characteristics) of EVs. Commonly used *in vitro* assays of EV bioactivity assess conditioned media (e.g. ELISA) or cell extracts (western blot, qPCR) from a target cell of interest in response to EV treatment, and essentially provide a population level assessment (22,23). Moreover, some MSC-EV bioactivity assays like wound healing and endothelial tube formation assays are helpful for understanding regeneration, but are not specific to a tissue or disease of interest (22,24). The performance of MSC-EVs in those assays may not be indicative of their function in a more tissue-specific disease context such as neuroinflammation. Thus, there is need for developing disease-specific tools to evaluate the bioactivity of MSC-EVs with a focus on treating neuroinflammation associated with chronic neurodegenerative diseases such as Alzheimer’s Disease (AD) or Parkinson’s Disease (PD).

Microglia play a key role in the onset and progression of neuroinflammation and have been explored as a therapeutic target for small molecule drugs and, more recently, cell therapies such as MSCs and MSC-EVs (25–28). As resident immune cells of the brain, microglia have been widely studied because of their role in homeostasis and disease pathology of the central nervous system (CNS) (29). Synaptic pruning, phagocytosis, remyelination and maintenance of vasculature are some of their key functions (26). Furthermore, they are phenotypically plastic, which is reflected by their morphological response to different microenvironmental cues (e.g. inflammatory cytokines). For instance, in patients that have suffered a stroke or have AD, there is a higher density of microglia that possess reduced number of processes and an increased cell body size (30,31). Studies have shown MSC-EV can modulate microglial secretion of pro-inflammatory cytokines, like IL-6, IL-1β and TNF-α, which are involved with neuroinflammation (32). Nevertheless, microglia exhibit significant heterogeneity at the single cell level and thus changes in production of individual secreted factors (or a limited panel) may be insufficient to fully represent the complex mechanisms of action of MSC-EVs (33–35). There have also been no studies demonstrating the effects of MSC-EVs on human microglia, as the entirety of the literature has used primarily non-human cell-lines, like primary rat/mouse cells, immortalized BV2 and N9 cell lines, among others (32,36,37). It is important to consider species-dependent differences in microglial response to MSC-EVs, as these differences could provide misleading interpretations of MSC-EV mechanism of action especially when *in vivo* animal models of neurodegenerative diseases do not fully capture key aspects of their analogous human disease (38,39). Moreover, despite the strong evidence that their morphology is associated with function (33,34), morphological profiling has not been applied to elucidate the bioactivity of cell-based therapies in the context of neuroinflammation.

Inspired from previous work (40–42), we developed a high-throughput, single-cell based microglia morphology assay as a readout of MSC-EV bioactivity that could inform their therapeutic potential. We comprehensively profiled the morphological response of an immortalized human microglia cell-line (43) upon stimulation with pro-inflammatory cytokines that are involved in neuroinflammation: IFN-γ (interferon-gamma) and TNF-α (tumor necrosis factor-alpha) (44–46). The activated microglia were then treated with MSC-EVs to assess the overall change in morphology. Phenotypic assessment of microglia was carried out using proteomic and lipidomic profiling following cytokine stimulation and upon MSC-EV treatment. This comprehensive approach of single cell morphological analysis combined with tools reflecting overall microglial phenotypic changes enabled us to demonstrate morphology as a robust, high-throughput outcome indicative of MSC-EV bioactivity. This assay can be further employed to screen different MSC cell-lines, EV manufacturing conditions and potentially elucidate their mechanism of action in the context of neuroinflammation.

## 2. Methods

### 2.1 MSC-EV manufacturing

A vial of cryopreserved, previously expanded, bone marrow-derived MSCs (donor RB71, additional donor information in **Table S1**) was thawed in RoosterBio growth medium (RGM) (RoosterNourish-MSC-XF [KT-016] containing RoosterBasal™-MSC (SU-005/SU-022) + RoosterBooster™-MSC-XF (SU-016)) on Day 0. On Day 3, ∼90% confluency, cells were split into 6 T-175 flasks (Greiner Bio-One, Cat#660175) at a seeding density of 1×10^6^ cells/flask in RGM. The media was supplemented with RB replenish (RoosterReplenish-MSC-XF [SU-023]) 24 hours post cell seeding (Day4). At ∼90% confluency (Day 6), spent media was aspirated and cells were washed once with sterile PBS-/-(Corning, Cat#21-040-CM) followed by addition of Rooster EV Collect media (RoosterCollect™-EV (M2001)) (Day7). The conditioned media was collected after 48 hours (Day9) for MSC-EV isolation.

MSC-EVs were isolated using a modified 2-step ultracentrifugation method (47). The media collected from 3 flasks (25mL/T-175flask) was filtered through a 0.2μm filter (VWR, Cat#10040-464) followed by first round of ultracentrifugation (Thermo Scientific™ Sorvall™ WX ultraCentrifuges, Rotor = F37L-8x100 −6202741) at RCFmax = 133,900 x g, for 1 hour @ 4℃. Each ultracentrifugation tube contains a maximum of 65mL spent media. The supernatant from this spin was discarded and the pellet was resuspended in cold PBS+/+ (Corning, Cat#MT21030CM) (4mL/tube) and collected in microultracentrifuge tubes and spun at RCFmax = 140,000 x g for 1 hour @ 4℃. The supernatant is discarded, and the pellet is resuspended in a total volume of 2mL PBS+/+. To mitigate any potential loss of MSC-EV bioactivity due to long-term storage and ensure consistency across all experiments, we used freshly prepared (i.e. never frozen) MSC-EVs for all experiments (48–50).

### 2.2 MSC-EV characterization

#### 2.2.1 Protein: Micro-BCA assay

The micro-BCA assay (ThermoFisher, Ca#23235) is used to quantify the amount of protein in our MSC-EV batch. We quantify the protein using 1X, 2X and 4X dilutions of the MSC-EV batch across different EV batches to ensure consistent manufacturing. 75μL of sample was combined with 75μL of the reagent mixture, uniformly mixed, and incubated for 2 hours in 37℃ (higher dilutions are diluted in PBS++ to match the 1X sample volume). The 96-well plate is read in a plate reader @562nm to quantify absorbance.

#### 2.2.2 Size, Count and Surface Marker: NanoFCM

MSC-EV size, concentration and tetraspanin surface marker expression (CD63) were assessed using the Flow Nanoanalyzer (NanoFCM, Inc., Xiamen, China). 1:100 dilution of MSC-EVs was used for assessment of MSC-EV size and concentration. MSC-EVs were stained with 20μM CFSE (BioLegend, Cat#423801) for 30mins at 37°C to indicate the presence of an intact lipid bilayer. FITC (10/50mW, 488) channel was used to assess %positive staining. CD63 (1:2000, BDBiosciences, Cat#561983) antibody was incubated with MSC-EV samples for 2 hours at room temperature (RT) prior to assessment. Surface marker intensity was measured using the APC channel (20/100mW, 638). Representative flow gating strategy for EV size, count, CFSE, and CD63 assessment can be found in **Figure S1**.

### 2.3 Microglia cell-line

C20 cells, an immortalized human cell-line, were generously provided by the Kim lab from Kent State University (Dyne et al., 2022;). The growth media consists of DMEM-F12 1:1 (Gibco, Cat#11320033), 10% FBS (Neuromics, Cat#FBS008, Lot#218H19), 1% PenStrep (Gibco, Cat#15070-063) and 0.2% normocin (InvivoGen, Cat#ant-nr-1). C20s are thawed in growth media containing 1% N2 supplement (Gibco, Cat#17502048) additionally for efficient recovery + 1 μM dexamethasone (Millipore sigma, Cat#D4902). The cells are recovered post thaw for 48 hours, reaching ∼90% confluence, followed by harvesting and seeding for experiments.

### 2.4 Microglia morphology assay

For morphology assays, cells are seeded in a Poly-D lysine hydrobromide (PDL) (Sigma-Aldrich, Cat#P6407) coated 96-well plate (Corning, Cat#3603; Greiner Bio-One, Cat #655090) at a density of 480 cells/well (1500cells/cm^2^) with 100μL of media/well. PDL coating aids in better adhesion of the cells preventing cell loss due to washing during staining. Post 24 hours of seeding, cells are stimulated (+CTL) using half media change with a cytokine cocktail of 5 ng/mL IFN-ү (Gibco, Cat# PHC4033) + 5 ng/mL TNF-α (Sino Biological, Cat#10602-HNAE). Treatment groups receive an additional 10 µL of MSC-EV preparation (+CTL+EVs). After 24 hours, cells are fixed, stained and imaged.

#### 2.4.1 Cell Staining and Imaging

The morphology plates are stained using a modified version of the Cell Painting protocol (53). The mitochondria are stained in live cells using Mitotracker Deep Red (Invitrogen, Cat#M22426) @ 10 ng/mL concentration for 30 minutes at 37℃. Cells are then fixed using 4% PFA (Electron Microscopy Sciences, Cat#15714-S) for 30 minutes and washed 2X with PBS-/- (Corning, Cat#21-040-CM) or 1X HBSS (Gibco, Cat#14065056). Further staining is carried out using HBSS 1X + 1%(w/v) Bovine Serum Albumin (BSA) (Sigma-Aldrich, Cat#A9418) + 0.1% Triton filtered through a 0.22μm filter. Wheat Germ Agglutinin, Alexa Fluor™ 555 Conjugate (WGA, 1.5μg/mL) (Invitrogen, Cat#W32464) Phalloidin/AlexaFluor 568 conjugate (8.25nM) (Invitrogen, A12380) and Hoechst(10ug/mL) (Invitrogen, Cat#H3570) are added for 30 minutes protected from light. The wells are washed three times with 1X HBSS and imaged using a 10X objective with a BioTek Cytation5 (Agilent) automated microscope. A 6 by 6 montage (36 total images) was captured for each replicate well with 6-8 technical replicate wells used for both control and EV treatment groups in each experiment.

#### 2.4.2 Image analysis

Single cell analysis morphological profiling was carried out with CellProfiler (54) using a custom pipeline (**Supplemental File F1**) The morphological data are passed through a series of processing steps. First, we combine all the channel files for cell, cytoplasm and nucleus resulting in a larger merged dataset. The experimental conditions are then added to the merged file which is further processed to add features like aspect ratio (major: minor axis length) and perimeter: area ratio. Poorly segmented outliers and/or debris are eliminated using cut off parameters like solidity=1 and nucleus: cytoplasm ratio>=0.85. Following image preprocessing, medians are calculated per well from the single cell data consisting of thousands of cells to ensure any other outlier effects are minimized. This median data is further normalized using min-max normalization where the average of negative controls(-CTL) is used as the min value and the average of positive controls (+CTL) represented the max value (**Supplemental File F2**). We then plot different features as well as perform principal component analysis (PCA) (55,56) on 21 cellular and nuclear features to assess the overall changes in microglia morphology.

### 2.5 Microglia secretion profile assessment

For secretion assays, C20 cells are seeded at a higher density of 6000 cells/well (18,750 cells/cm^2^) in a 96-well plate with 100μL of media. The stimulation and treatment strategy are identical to the morphology assay (concurrent cytokine and EV treatment) followed by collection of the spent media from each well after 24 hours. The collected media is stored at −80℃ and then shipped to RayBiotech Inc. on dry ice for screening a panel of 200 targets (Human Cytokine Array Q4000, QAH-CAA-4000).

### 2.6 Microglia Lipidomic profile assessment

C20 cells were seeded at a density of 100,000 cells/cm^2^ in a 6-well plate in 2mL of media. The timeline and treatment procedure was the same as morphology experiments with a half media change for stimulation and treatment carried out 24 hours post seeding. 24 hours post treatment, cells were washed 3X with 1mL cold sterile PBS-/- (Corning, Cat#21-040-CM) and harvested with 300μL of TrypLE (Gibco, 12604-013) for 5 mins in 37℃, neutralized with 1.5mL of growth media and spun at 490g for 7 minutes. The supernatant was aspirated and the pellet was resuspended in 200μL of extraction solvent (80% methanol solution (MeOH)). The suspension was then flash frozen in liquid nitrogen and stored in −80℃ freezer for further processing.

Lipids were obtained from cell samples using a biphasic extraction method and identified using LC-IM-MS/MS. Sample preparation for LC-MS was carried out using the methods from Hewelt-Belka W et. al. (57). In brief, pellets were resuspended in 0.5-1mL of water or 4:1 MeOH /H20, vortexed briefly and transferred to a 5mL glass centrifuge tube. Blank was prepared similarly. All samples were sonicated and placed in an ice bath for 30mins followed by addition of 2mL 1:2 chilled chloroform/methanol solution. The samples were then vortexed on and off for 5min with storage on ice between vortex cycles. 0.5mL of chilled chloroform followed by 0.5mL of chilled water was added to the samples and vortexed for 60seconds. The samples were centrifuged for 10mins at 4°C and 3,500g. The bottom organic layer was transferred to a new 5mL glass tube followed by drying in a vacuum concentrator (Savant, ThermoScientific). Dried extracts were reconstituted with 500μL of 1:1 chloroform/methanol solution. Using pasteur pipettes, extracts were transferred to 2mL glass vials and stored at −80°C. For mass spectrometry sample preparation, each extract was prepared as a 2.5X dilution in HILIC A (95:5 ACN/H_2_0, 10mM ammonium acetate), with 10μL of each sample combined for QC. Mass spectrometry was carried out using the Waters Synapt XS instrument with a 100mm BEH-HILIC column with HILIC A (95:5 ACN/H2O, 10 mM Ammonium Acetate) and HILIC B (50:50 ACN/H2O, 10 mM Ammonium Acetate) used as mobile phases (58). Samples were collected for 7 minutes. Data were collected using MassLynx and processed with Progenesis QI, EZInfo, and Lipostar software. Lipid pathway enrichment analysis was carried out using LIPEA (59).

### 2.7 Microglia Cellular Proteomics Analysis

C20 cells were seeded at a density of 10^5^ cells/well in a 6-well plate in 2mL of media. The timeline and treatment procedure were the same as morphology experiments with a half media change for stimulation and treatment carried out 24 hours post seeding. 24 hours post treatment, cells were harvested with 300μL of TryplE (Gibco, 12604-013) for 5 mins in 37℃, neutralized with 1.5mL of growth media and spun at 490g for 7 minutes. The pellets were resuspended in sterile PBS-/- (Corning, Cat#21-040-CM) for a wash followed by the spin settings mentioned earlier. The supernatant was aspirated and cell pellets were flash frozen in liquid N2 and stored in −80C. The tubes were then transported to the core facility on dry ice.

Approximately half a million cells were lysed in a lysis buffer containing 5% SDS, 50 mM TEAB (pH 8.5) and 1X protease and phosphatase inhibitor cocktail and MgCl2 (2mM) at 95°C for 15 minutes. The lysate was incubated in 100 units of benzonase for 5 minutes at room temperature. The lysate was centrifuged at 21000g for 10 minutes and the supernatant was collected. Protein concentration was estimated by nanodrop and 50 micrograms of protein was used for tryptic digestion. Protein concentration was adjusted to contain 50 micrograms in 23 microliters of lysis buffer and 1microliter of 100 mM TCEP was added to the lysate and incubated at 55°C for 15 minutes. Sample was alkylated by adding 1 microliter of Iodoacetamide (200 mM) and incubating in dark at room temperature for 15 minutes. Sample was acidified by adding 2.5 microliters of phosphoric acid (55%) to the tube. The sample was transferred to S-traps after mixing it with 165 microliters of S-trap binding/wash buffer (90% methanol in 100 mM TEAB). S-trap column was placed in a collection tube and centrifuged at 4000g for 1 minute to bind the protein to the resin. Protein was washed thrice with 150 microliters of S-trap binding/wash buffer. The proteins were digested by adding 2.5 micrograms (1:20) of trypsin in 20 microliters of 50mM TEAB to the trap and incubating it at 37°C for overnight. Tryptic peptides were eluted by adding 40 microliters of elution buffer1 (50 mM TEAB) and spinning the trap at 4000g. Elution was repeated with elution buffer 2 (0.1% Formic Acid) and elution buffer 3 (50% Acetonitrile). The pooled eluate was dried in speedvac. Samples were then reconstituted in 20 µL of 0.1% formic acid (FA). To facilitate peptide resuspension tubes were incubated in a thermomixer at 2000 rpm and 37°C for 10 minutes and centrifuged at 21000 g for 10 minutes at room temperature. Ten microliters of digested peptides were carefully transferred from the top of the microfuge tube to the sample vial. A one microgram equivalent of peptide was subjected to LCMS/MS analysis on Orbitrap Q-Exactive Plus mass spectrometer (Thermo Scientific) in conjunction with Dionex UltiMate3000 Ultra High-Performance Liquid Chromatography (Thermo Scientific) system. An externally calibrated Thermo Q Exactive Plus (high-resolution electrospray tandem mass spectrometer) was used in conjunction with Dionex UltiMate3000 RSLCnano System. A 2 μL sample was aspirated into a 50 μL sample loop and loaded onto the trap column (Thermo µ-Precolumn 5 mm, with nanoViper tubing 30 µm i.d. × 10cm). The flow rate was set to 300 nL/min for separation on the analytical column (Acclaim pepmap RSLC 75 μM× 15 cm nanoviper). Mobile phase A was composed of 99.9% H2O (EMD Omni Solvent), and 0.1% formic acid and mobile phase B was composed of 99.9% ACN, and 0.1% formic acid. A 120-minute step gradient from 3% to 45% B was performed. The LC eluent was directly nanosprayed into Q Exactive Plus mass spectrometer (Thermo Scientific).During the chromatographic separation, the Q Exactive plus was operated in a data-dependent mode and under direct control of the Thermo Excalibur 3.1.66 (Thermo Scientific). The MS data were acquired using the following parameters: 20 data-dependent collisional-induced-dissociation (CID) MS/MS scans per full scan (400 to 1800 m/z) at 70,000 resolution. MS2 were acquired at 17,500 resolution. Ions with single charge or charges more than 7 as well as unassigned charge were excluded. A 10 second dynamic exclusion window was used. All measurements were performed at room temperature. Resultant Raw files were searched with Proteome Discoverer 3.0 using SequestHT as the search engine using human proteome FASTA database downloaded from Uniprot. Percolator was used as peptide validator.

#### 2.7.1 Gene set variation analysis

To study functional impact of proteomic changes within each experimental group, gene set variation analysis (GSVA) was conducted. GSVA is an unsupervised enrichment algorithm which identifies variations of pathway activity by defining enrichment score for gene sets which each contain a set of genes that share same cellular function. To enable GSVA, we used the Molecular Signatures Database C2 [v7] gene sets (MSigDB) (60), which represents cell states and perturbations within the immune system. The *gsva* R package (available on Bioconductor) (61) was used to compute enrichment of immune related signatures within each group. To evaluate statistical differences between groups for each pathway of interest, we employed ANOVA followed by Tukey post-hoc test, then Benjamini–Hochberg false discovery rate (FDR) adjustment for multiple comparisons were used. FDR adjusted p-values≤0.05 were considered significant (**Supplemental File F3**).

#### 2.7.2 Hierarchical clustering for heatmap

Data were z-scored for each protein across samples and clustered using the Euclidian distance metric and the Ward. D2 agglomeration method.

#### 2.7.3 Differential abundance analysis

Differential abundance analysis was conducted using the *limma* package in R (62). Treatment was used as the variable in the design matrix. Significance level (p-value, linear model) and fold changes were computed using the *lmFit()* function. |Fold Change|≥2 and p<0.05 were considered to be significant (**Supplemental File F4**). Differentially Abundant Proteins (DAPs) were displayed as volcano plots using *EnhancedVolcano* R package (63), available through Bioconductor.

#### 2.7.4 Gene Ontology enrichment analysis

Gene Ontology (GO)_was conducted for differentially abundant proteins using the PANTHER overrepresentation test with PANTHER 18.0 through the Gene Ontology resource (64). The *homo sapiens* GO biological processes complete annotation set was used with Fisher’s exact test, followed by Benjamini–Hochberg false discovery rate (FDR) adjustment to compute significance of the biological processes linked with differential abundant proteins. The reference list consisted of all detectable proteins in our dataset after filtering.

### 2.8 Statistical Analysis

Graphpad Prism 9 was used to perform all univariate statistical tests. Each figure legend describes the tests used for the respective experiment.

## 3. Results

### Microglia increase in size and complexity upon stimulation with cytokines

Upon stimulation with IFN-ү and TNF-α (5 ng/mL each, +CTL), we observed significant changes in microglia morphology (**Figure 1**), including perimeter, form factor and major axis length (**Figure 1B)**. Increase in perimeter and major axis length illustrate the characteristic increased cell size and elongation of +CTL cells compared to -CTL (unstimulated) cells. +CTL cells were more ‘complex’ as indicated by reduced form factor, a measure of the roundness of the object (form factor for circle=1) (54).

**Figure 1:**
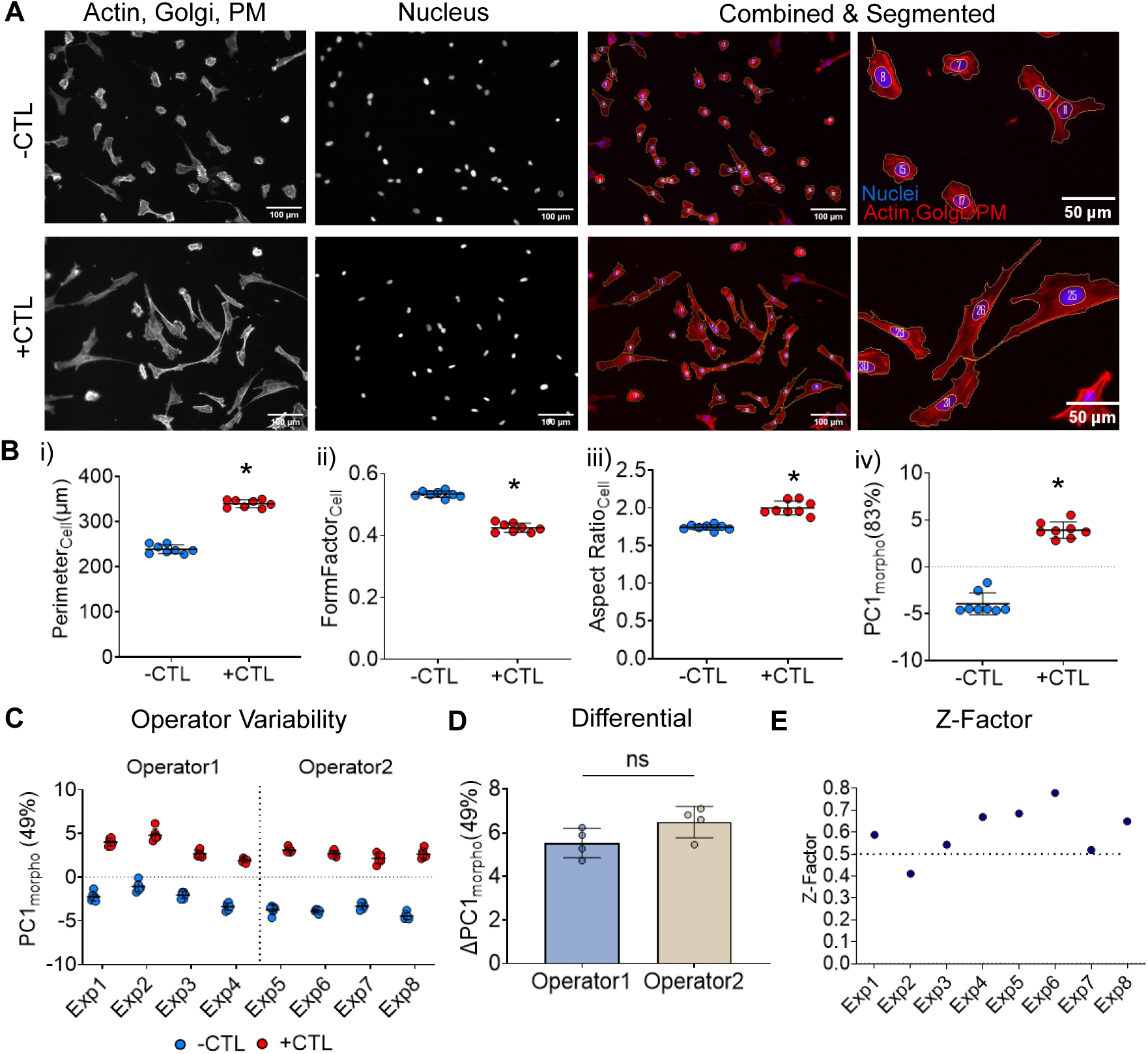
Microglia morphology changes upon activation with cytokines. **(A)** Representative images of C20 microglia in -CTL group (unstimulated) and +CTL group (stimulated with 5ng/mL each of IFN-γ and TNF-α). Actin, Golgi & plasma membrane (PM) (Red, Phalloidin/WGA), Nuclei (Blue, Hoechst): Scale bar=100μm **(B)** Individual cellular features (i) Perimeter, (ii) Form Factor and (iii) Aspect Ratio and (iv) Principal component 1 (PC1_morpho_) calculated using 21 features. Each point is a median of ∼700-1000 cells/Fig 1B well with mean and standard deviation plotted for n=8 wells per group. *p<0.01 vs -CTL for all groups conducted using unpaired t-test with Welch’s correction. **(C, D)** Experimental and Operator variability: Microglia controls (-CTL vs +CTL) are significantly different across multiple experiments and operators. p<0.001 vs +CTL for all experiments using 2-way ANOVA with Šídák’s multiple comparisons test. **(E)** Z-factor for PC1_morpho_ is > 0.5 (dotted line) for 7/8 experiments.

To better account for the contribution of each feature to the overall morphology (and considering that a single feature does not fully encompass each microglia’s morphological response), we used PCA to effectively create a composite overall morphological score, as previously shown using MSCs (40,41). The first principal component (PC1_morpho_) accounted for 83% of the variance in the dataset and thus was used to represent an overall composite morphological response of microglia (**Figure 1B(iv)**). This trend across individual parameters as well as PC1_morpho_ is consistent across multiple experiments (n=5) as shown in supplementary **Figure S2.**

### Demonstration of morphological assay reproducibility and robustness

We quantified the reproducibility and robustness of our morphological assay based on experiment-experiment variability and different assay operators, respectively. In **Figure 1C**, we showed a consistent change in overall microglia morphology between operators O1 and O2 for 4 independent experiments. Despite some observed experimental variability within a given operator (data was normalized to controls within an experiment but not between experiments), the differential morphological response (indicated by ΔPC1_morpho_, **Figure 1D**) is not significantly different between experiments or operators. Moreover, for high throughput screening assays, a common metric of assay quality is the Z-factor (Z’) (65). Used extensively in drug screening, it is calculated using the following formula:

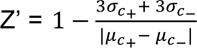

Where,

σ= standard deviation

c+ = positive control (+CTL)

c- = negative control (-CTL)

μ = mean

Across all but 1 of our experiments (**Figure 1E**), the Z-factor calculated using PC1_morpho_ values is > 0.5. Based on Z’ value classification, an assay with a Z’> 0.5 corresponds to an optimized assay with a desirable dynamic range (65).

### Microglial lipid content increases upon stimulation

Nine different classes of lipids were identified (**Figure 2A**), including phosphatidylcholines (PCs), phosphatidylethanolamine (PE), phosphatidylglycerols (PGs), plasmalogen phosphatidylethanolamines (PE-Ps), sphingomyelins (SMs), Ceramides (Cer), Hexosylceramides (HexCer), triglycerides (TGs) and lysophosphatidylcholines (LPCs). Additionally, a higher number of individual lipids were detected for PCs, PEs and PGs compared to other lipid classes. An increase in all lipid classes except LPCs was observed upon microglia stimulation, which may be attributed to the increased cell size. Increased lipid accumulation has been linked to pro-inflammatory microglia with compromised ability to perform critical functions like phagocytosis (66,67). Lipid pathway enrichment analysis (LIPEA) (59) was used to identify the various pathways associated with the increased lipid content. Ferroptosis and sphingolipid pathways, which have been associated with neurodegeneration (68–70), were enriched post microglia stimulation (**Figure 2B**).

**Figure 2:**
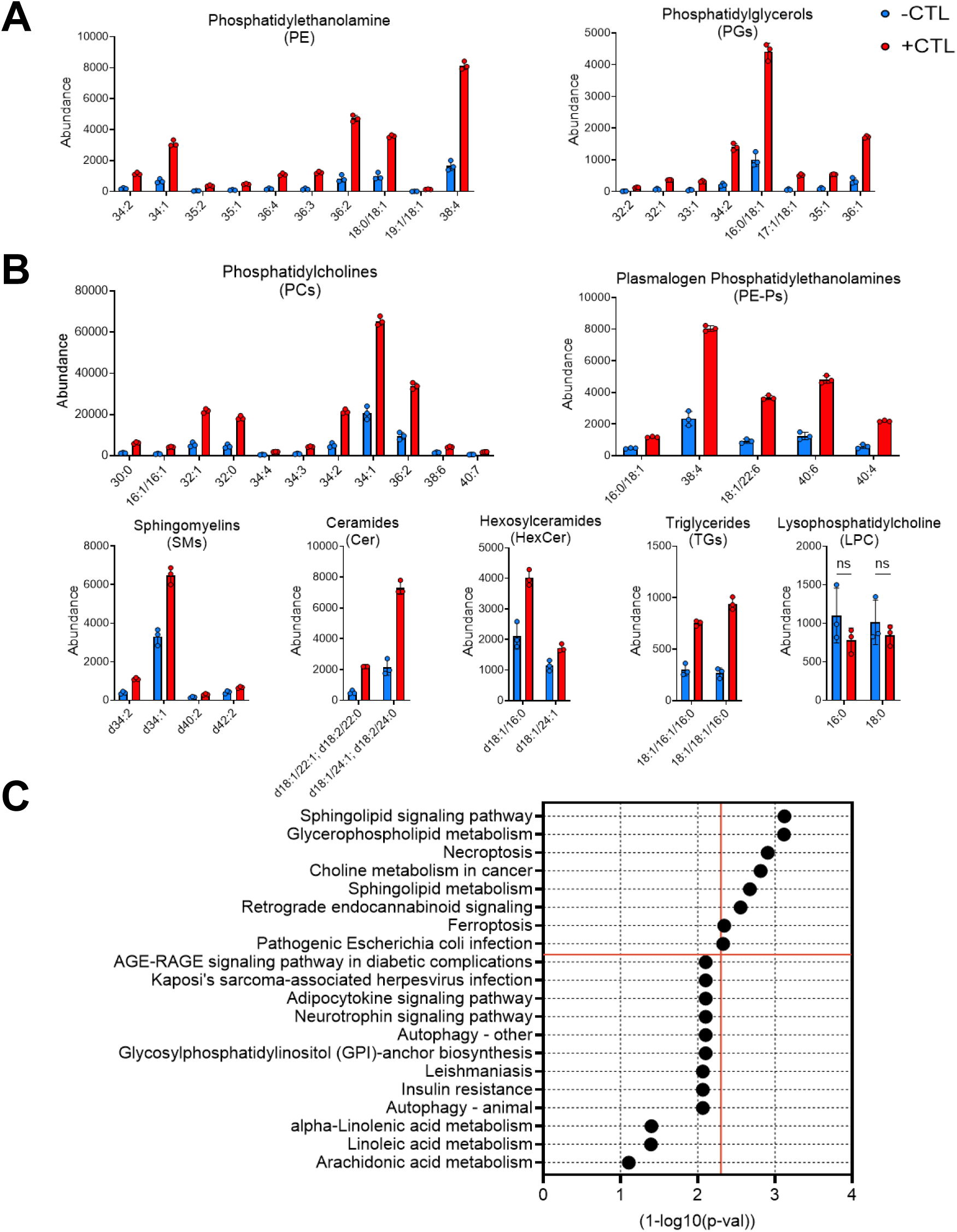
Microglia lipid content increases upon activation with cytokines. **(A)** Overall cellular lipid content across different lipid classes increases following stimulation (+CTL). Lipid classes identified using negative mode **(A)** and positive mode **(B)** ionization. Each lipid species significantly different between –CTL and +CTL (p<0.05) except LPC (denoted by ‘ns’) using unpaired t-test with Welch’s correction. **(C)** Pathways ordered by p-value significance (1-log10(p-val)) with significantly enriched pathways above red horizontal line at p-value cutoff = 0.05.

### Activated microglia secrete factors that relate morphology with inflammation

Across both unstimulated (-CTL) and stimulated (+CTL) microglia, we detected quantifiable secretion of 94 cytokines, chemokines and growth factors from a 200-plex antibody array panel (**Figure 3A**). Of the 94 targets (**Table S2**), 40 of them were significantly different between controls with increased levels of 38 proteins and decreased levels of only 2 proteins following cytokine treatment. We conducted pathway analysis using the 40 significantly different proteins as an input to the STRING database (**Figure 3B**). The map shows the physical and functional interactions between all the proteins (nodes) that were increased (red) and decreased (green) upon stimulation. An overview of proteins differentially secreted between −/+CTL groups ordered in terms of the number of connected node degrees with other proteins in this network (**Table S3**) is shown in **Figure 3C**. Notably, the cytokines/chemokines IL-6, CXCL8 and CCL2 had the highest number of connections and were secreted at much higher levels in +CTL groups vs. -CTL microglia.

**Figure 3:**
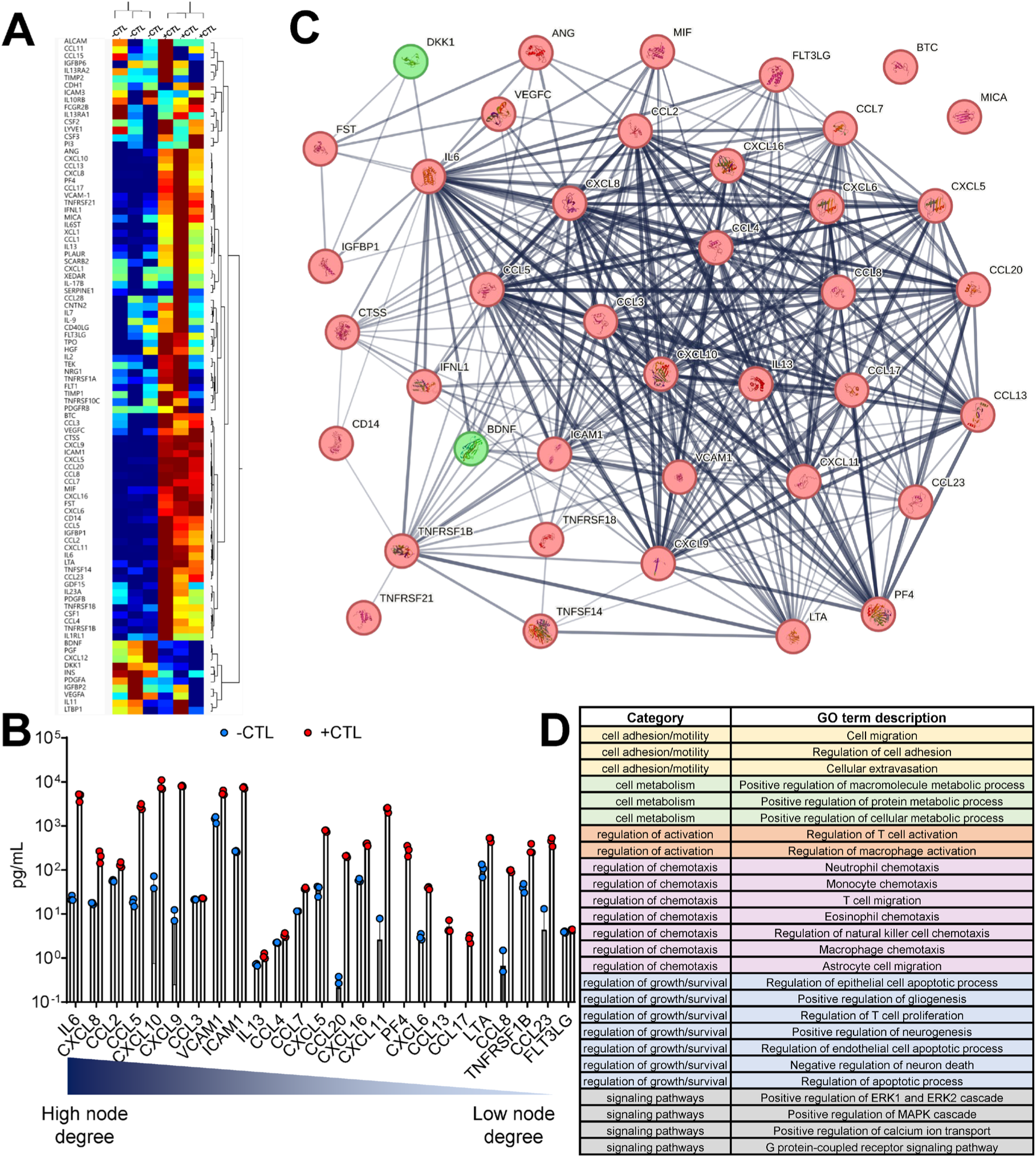
Effects of cytokine stimulation on microglia secretion. **(A)** Heatmap representing microglia proteins that change following IFN-γ/TNF-α stimulation (-CTL vs +CTL). Two-way hierarchical clustering using Ward method. **(B)** STRING database generated network map of proteins increased (red) and decreased (green) after stimulation (+CTL) and their predicted functional associations (edges). Thicker and darker edges (line connecting 2 proteins), implies higher level of confidence in the interactions and correlation between the proteins (http://version10.string-db.org/help/faq/). **(C)** Proteins that are significantly different between controls and arranged in descending order of their number of functional associations (node degrees). p<0.05 using unpaired t-test with Welch correction for all proteins. **(D)** Pathways associated with changes in microglia secretion post stimulation.

Using the entire proteome as a background, we performed pathway enrichment analysis (based on Gene Ontology (GO) Terms) using STRING to identify potential disease-relevant pathways of microglia associated with changes in secretion and morphology. The entire list of pathways is available in **Table S4** with a curated subset of pathways of interest shown in **Figure 3D**. These pathways were selected based on categories relevant to observed morphological response (i.e. cell adhesion/motility), cell metabolism, regulation and activation of cells, as well as specific signaling pathways. Of note, many of these pathways involve microglia regulation of other cells involved in neuroinflammation such as T cells, macrophages, and astrocytes through processes such as chemotaxis and growth/survival (71,72). We also performed enrichment analysis using our 200plex panel as a background, but no pathways were significantly enriched; however, this is not unexpected as this a preselected, focused panel of chemokines/cytokines/growth factors that are commonly produced by microglia both *in vitro* and *in vivo*.

### MSC-EV treatment modulates microglia morphological response to cytokines

MSC-EV treatment prevents the morphological shift of stimulated microglia by essentially inhibiting this cytokine-induced change in morphology of increased size/complexity (**Figure 4A**). Morphological features were normalized based on controls (using max-min approach) to reflect the relative morphological change after MSC-EV treatment. With concurrent MSC-EV treatment, the microglia are effectively smaller and less complex compared to +CTL microglia as reflected by differences in the normalized features like perimeter, aspect ratio and form factor, respectively (**Figure 4B**). The overall change in morphology assessed using composite morphological score PC1_morpho_ (72.2% variance) indicates a shift in the morphology of MSC-EV treated microglia towards the -CTL microglia. This trend was observed across multiple experiments (using independently manufactured MSC-EV batches from the same cell-line to ensure consistency), as shown in **Figure S3**.

**Figure 4:**
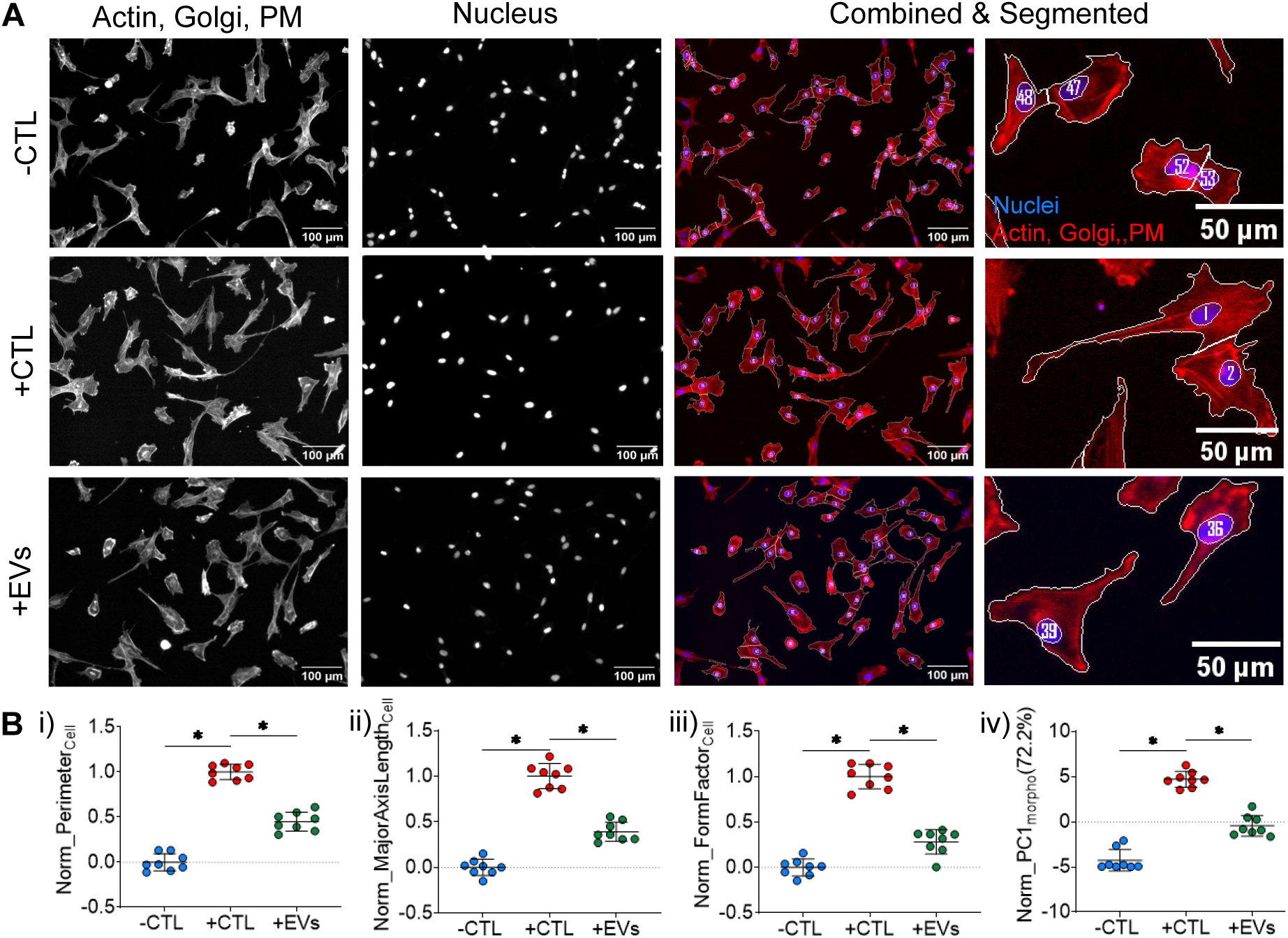
Stimulated microglia morphology changes upon treatment with MSC-EVs. **(A)** Representative images of C20 microglia cells in -CTL group (unstimulated), +CTL group (stimulated with 5ng/mL each of IFN-γ and TNF-α) and +CTL+EVs (stimulated with 5ng/mL each of IFN-γ and TNF-α + treated with MSC-EVs). Actin, Golgi & plasma membrane (PM) (Red, Phalloidin/WGA), Nuclei (Blue, Hoechst), Scale bars =100 μm (columns 1-3 images) and 50 μm (column 4 images). **(B)** Individual min-max normalized cellular features **(i)** Perimeter, **(ii)** Major Axis Length and **(iii)** Form Factor. **(iv)**Normalized principal component 1(PC1_morpho_) calculated using 21 min-max normalized features represents 72.2% of the variance. Each point is a median of ∼700-1000 cells/well. Bar indicates the mean of the 8 wells/group. *p<0.001 vs +CTL for all groups conducted using Brown-Forsythe and Welch one-way ANOVA with multiple corrections using Dunnett T3 testing.

### Microglial lipid content does not change in response to MSC-EVs

Despite the changes in morphology, no changes in lipid content were observed in response to MSC-EV treatment (**Figure 5**). The detected amounts were unaltered between different classes of lipids across the stimulated (+CTL) and the MSC-EV treated groups (+EVs). Therefore, no pathway enrichment analysis was carried out. However, the PCA plot shows overall changes in the lipid profiles (using lipid feature intensities instead of annotated lipids as input to PCA) between activated and MSC-EV treated cells (**Figure S4**) suggesting other unidentified/unannotated lipids may be different between the 2 groups.

**Figure 5:**
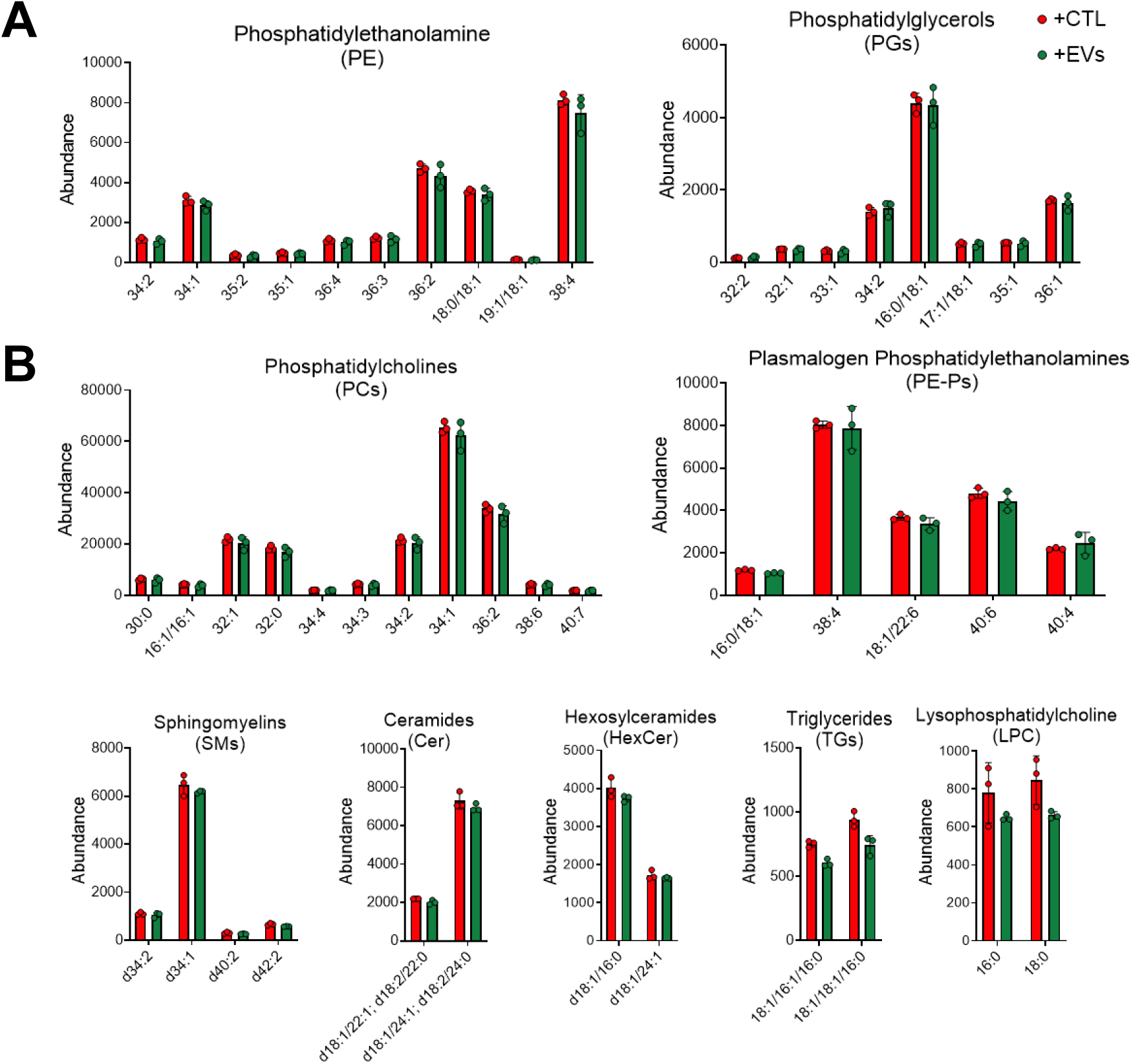
Microglia lipid content remains unchanged with MSC-EV treatment. Similar amount of lipids observed between the stimulated (+CTL) and MSC-EV treated (+EVs) microglia. Lipid classes identified using negative mode **(A)** and positive mode **(B)** ionization. P>0.05 vs +CTL for all lipids calculated using unpaired t-test with Welch’s correction.

### Mitochondrial morphology changes in response to MSC-EV treatment

Overall changes in mitochondrial morphology were assessed using PCA on 225 mitochondrial features across different treatment conditions (**Figure 6A**). PC1_mito_ captures the maximum variance in data (58.1%) using features including the intensity, distribution and texture of the mitochondria (stained using Mitotracker Deep Red, **Table S5**). In accordance with the cellular morphology, the mitochondrial morphology is also significantly different between unstimulated (-CTL) and stimulated (+CTL) microglia (*p<0.05, **Figure 6B,C**) across 4 independent experiments using 2 different cell culture plate-types. Additionally, MSC-EV treatment (+EVs) results in a greater significant change in mitochondrial morphology in 2 out of 4 experiments where notably the type of tissue culture plates may impact the significance of the observed changes with more consistent shift (i.e. decrease in PC1_mito_) when using plates with µClear® film bottom (**Figure 6C**).

**Figure 6:**
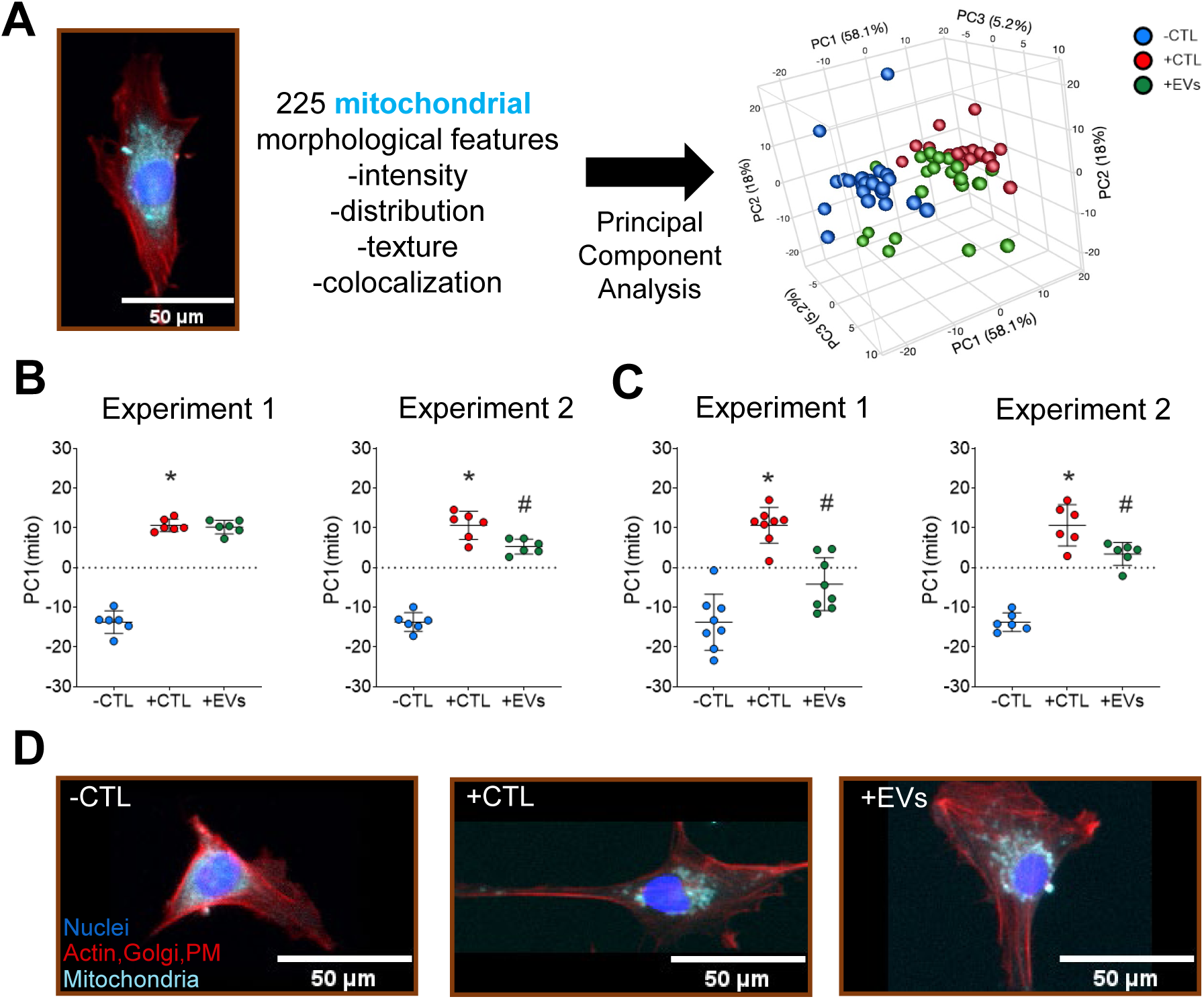
Microglia mitochondrial morphology altered by MSC-EV treatment. **(A)** Representative image of microglia mitochondria (Cyan) used to calculate single-cell features. Principal component analysis indicates maximum variance in PC1_mito_ (58.1% variance) **(B &C)** PC1_mito_ used as composite mitochondrial morphological score to assess effects of with (+CTL) and without (-CTL) cytokine stimulation and concurrent treatment with MSC-EVs (+EVs). Assay conducted on Costar Cat#3603 **(B)** and Greiner Cat#655090 **(C)** 96-well plates, respectively. *p<0.05 vs -CTL and #p<0.05 vs +CTL conducted using Brown-Forsythe and Welch one-way ANOVA with multiple corrections using Dunnett T3 test. **(D)** Representative images of mitochondria from each experimental group. Actin, Golgi & plasma membrane (PM) (Red, Phalloidin/WGA), Nuclei (Blue, Hoechst) and Mitochondria (Cyan, Mitotracker Deep Red), Scale: 50μm.

### MSC-EVs modulate the secretion profile of stimulated microglia

Using the same 200plex antibody array, we profiled microglia secretome in response to MSC-EV treatment. PCA (**Figure 7A**) was conducted using 24 proteins (**Figure 7B**) that were significantly different between +CTL and +EV groups. Distinct clusters were apparent for all 3 experimental groups using a plot of principal components PC1_secretion_ (51.8%) and PC2_secretion_ (36.5%) that accounted for the majority of data variance. Changes in overall secretome reflected by PC1_secretion_ suggest a different (‘alternate’) response of microglia to EVs as compared to the morphological, mitochondria, and lipidomic responses. Notably, PC1_secretion_ scores for the +EV group were significantly different from both -CTL and +CTL, but these controls were not different from each other. PC2 _secretion_ scores, on the other hand, exhibited the previously observed behavior of MSC-EVs inhibiting the microglia response (in this case secretion). **Figure 7C** shows the pathway analysis conducted using the STRING database on the subset of 24 proteins differentially regulated by MSC-EVs. Secretion of 14 proteins decreased (green nodes) and 10 proteins increased (red nodes) in response to MSC-EV treatment (**Figure 7C(i)**). These proteins were found to be linked with some of the previously identified metabolic and cell migration pathways in control groups (**Figure 7C(ii)**). Moreover, new functional pathways related to regulation of inflammation and EV-mediated responses (e.g. regulation of cytokine/chemokine production, regulation of angiogenesis) were identified. These differentially regulated proteins were also found to be involved in a number of non-microglia cell pathways such as T cell proliferation, neurogenesis, and immune cell migration (e.g. macrophages, natural killer cells and T cells). A complete list of all enriched pathways, including the differentially secreted proteins associated with each pathway, can be found in **Table S6**.

**Figure 7:**
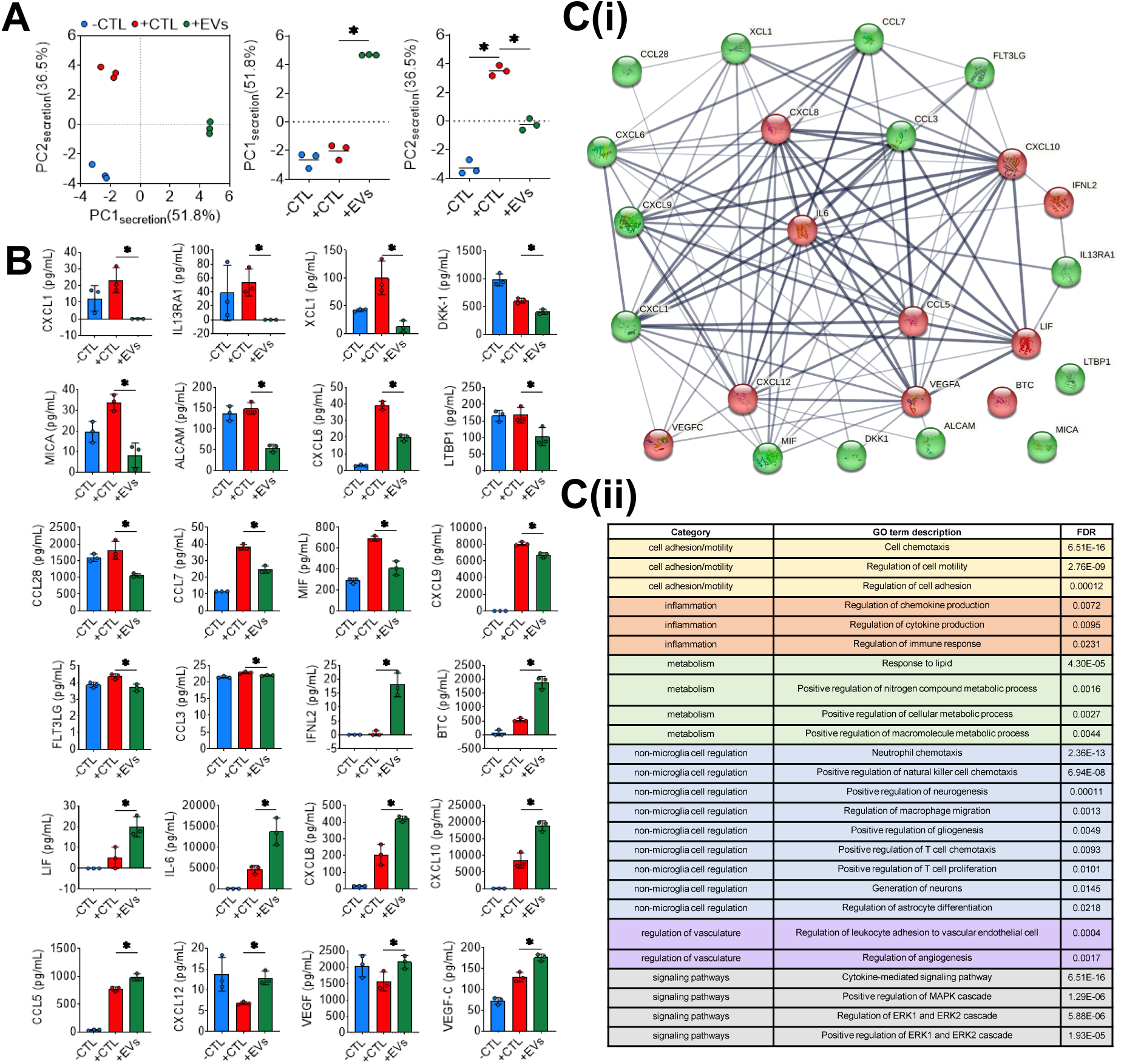
Microglia secretion profile is altered post MSC-EV treatment. **(A)** Individual clusters of -CTL (blue), +CTL (red) and +EVs (green) observed with PCA conducted using 24 proteins that were different between +CTL and +EVs. PC1_secretion_ (51.8% variance) and PC2_secretion_ (36.5% variance) account for 88.3% of total variance. **(B)** Observed changes in secretion of individual proteins (14 decreased, 10 increased) with concurrent MSC-EV treatment. *p<0.05 conducted using unpaired t test with Welch correction. **(C(i))** STRING database generated network map of for group of proteins decreased (green) and increased (red) after MSC-EV treatment and their predicted functional associations (edges). Thicker and darker edges (line connecting 2 proteins), implies higher level of confidence in the interactions and correlation between the proteins (http://version10.string-db.org/help/faq/). **(C(ii))** Pathways associated with changes in microglia secretion following MSC-EV treatment.

### Alterations in microglia proteome in response to MSC-EVs

Using LC-MS/MS, 2662 proteins were identified across the 3 experimental groups -CTL, +CTL and +EVs. **Figure 8A** is a heatmap representation of z-scored expression of 2662 proteins across three different experimental groups suggesting distinct proteomic profile across groups, clustered based on their Euclidian distance. PCA revealed separation of experimental groups into distinct clusters based on their overall cellular proteomic profile (**Figure 8B**). Like the overall secretion profile (PC1_secretion_, **Figure 7A**), PC1_proteome_ (34.2% variance) suggested an alternate proteomic profile for the MSC-EV treated microglia compared to both controls. PC2_proteome_ (26.9% variance) corresponded to shift in proteomic profile towards the -CTL in response to MSC-EVs similar to the overall morphological response. Of the 2662 proteins, we identified 687 differentially abundant proteins (DAP) where 258 proteins were upregulated and 429 proteins were downregulated in MSC-EV group (|Fold Change|≥2 vs. +CTL microglia, p<0.05, linear mixed models) (**Figure 8C**). The differential abundance analysis as represented using a volcano plot in **Figure 8C** highlights certain proteins that were upregulated (e.g. RHOC, RHOA, PAC2) and downregulated (e.g. RAB23, PTEN, ACP1) in MSC-EV treated microglia as compared to +CTL microglia (significance p<0.05 and |Fold Change|≥2 vs. +CTL microglia). All 2662 proteins were used to conduct gene set variation analysis (GSVA, **Methods**) to identify the pathways that were altered in response to MSC-EV treatment with a subset of 8 gene sets highlighted in **Figure 8D**. Some pathways (e.g. TGF-β, Platelet homeostasis) were enriched in the +CTL vs. -CTL microglia while others (e.g. TNFR1, Semaphorin) were suppressed. In some cases, MSC-EV treatment modulated the pathway expression towards the -CTL; however, for certain pathways (e.g. Rac1, IL-4 and IL-13 signaling) MSC-EVs had an alternate response compared to the control groups and were highly enriched. The complete list of pathways that were significantly altered can be found in **Table S7** (enriched gene sets) and **Table S8** (suppressed gene sets).

**Figure 8:**
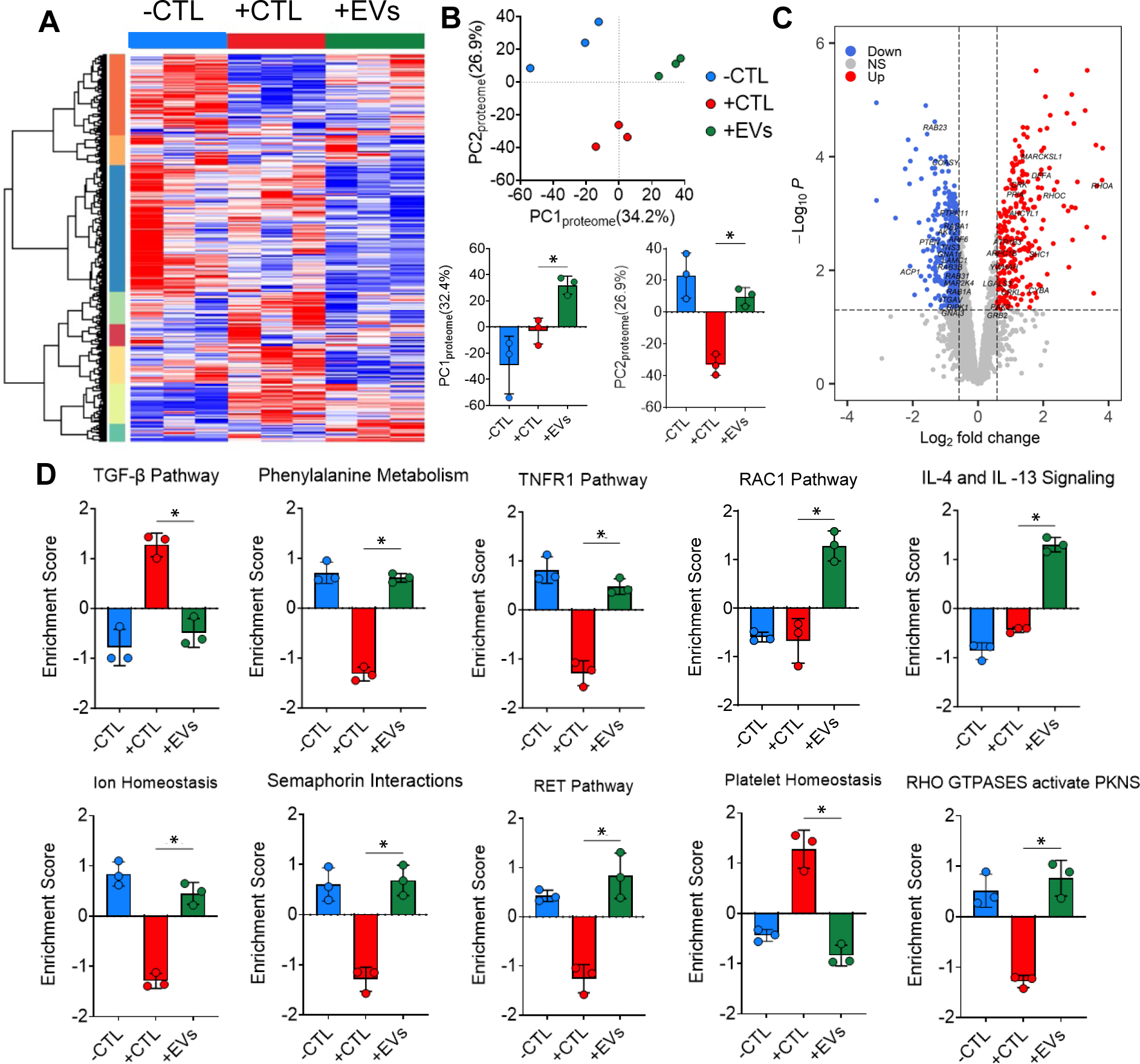
Microglia proteomic profile shifts after cytokine stimulation and with MSC-EV treatment. **(A)** Heatmap representing the 2662 proteins identified using LC-MS/MS and clustered based on Euclidian distance across all 3 experimental groups. **(B)** PCA reveals overall change in protein profile with PC1_proteome_ (34.2% variance) accounting for alternate protein expression and PC2_proteome_ (26.9% variance) accounting for shift in protein expression towards -CTL after MSC-EV treatment. **(C)** Volcano plot depicting alterations in protein expression upon MSC-EV treatment vs +CTL. **(D)** Subset of enriched pathways identified using GSVA analysis. *p<0.05 conducted using ANOVA followed by Tukey post-hoc test, then Benjamini–Hochberg false discovery rate (FDR) adjustment for multiple comparisons were used. FDR adjusted p-values≤0.05 were considered to be significant.

## 4. Discussion

In this study, we have shown that morphological profiling of microglia *in vitro* is a robust, reproducible approach to assess microglia phenotype relevant to neuroinflammation in response to a cell-derived therapeutic: MSC-EVs. We illustrated that significant changes in microglia morphology in response to IFN-ү and TNF-α correspond to increased production of other pro-inflammatory cytokines (e.g. IL6, CXCL8, CCL5, CCL2, CCL3), as well as increased lipid production (e.g. PCs, PEs), which are common indicators of microglia function (46,67,73–75). Additionally, pathways linked to metabolic activity (e.g. sphingolipid metabolism), cell motility (e.g. migration, extravasation) and regulation of immune cells (e.g. macrophages, T cells) are enriched upon exposure to cytokines and have previously been linked to inflammatory microglia profiles (76–79). With consistent controls and reproducible results across multiple experiments and operators, this assay enables high throughput, single-cell profiling of microglia in the context of neuroinflammation.

Using this assay, we were able to demonstrate that microglial morphological response to MSC-EV treatment is indicative of a desired, therapeutic effect in terms of microglia secretion. Other studies using non-human microglia have shown that changes in morphology in response to a therapeutic treatment (e.g. dexamethasone) can reduce secretion of inflammatory cytokines (e.g. TNF-α, IL-1β) (80). Similar behavior was observed in our studies as microglia had significantly reduced levels of inflammatory cytokines in response to MSC-EVs (e.g. CXCL6, CXCL9, CCL7). Pathway analysis carried out using differentially regulated proteins suggest MSC-EVs may exert their effects on microglia through modulation of inflammation, metabolism, regulation of vasculature and specific signaling pathways like MAPK and ERK. Chemokines like CXCL9 and CCL3 are involved with recruitment and proliferation of T cells which may aid in regulation of the immune response and thus inflammation (81,82). However, it’s important to note that MSC-EV modulation of microglia does not only result in an overall reduction in secretion of chemokines/cytokines as we also observed an increase in several proteins such as CXCL8, CXCL10 and CCL5. While commonly associated with inflammation, CXCL8 and CXCL5 have also been shown to relay neuroprotective effects by regulating neuronal apoptosis (83,84). Increased levels of these cytokines with MSC-EV treatment may relay a similar effect as pathways like ‘positive regulation of neurogenesis’ and ‘generation of neurons’ were associated with the proteins differentially secreted in response to MSC-EVs (**Figure 7D**). These differences may influence the recruitment and activation of different immune cell sub-types like T-cells, NK cells and macrophages to the site of inflammation. Regulation of blood brain barrier (BBB) permeability is critical to promote immune cell recruitment, and dysregulation/breakdown of the BBB is a hallmark of many neurodegenerative diseases. Increased secretion of vascular endothelial growth factors (VEGFs), which occurs in ischemic stroke and intracerebral hemorrhage has been linked with improved BBB permeability (85–87). Moreover, treatment with VEGF and VEGF-C have been shown to improve microglia phagocytosis of amyloid beta oligomers (88) and induce an anti-inflammatory profile in rat traumatic brain injury model (89). By producing these cytokines in the brain, microglia also promote angiogenesis, neurogenesis and facilitate indirect secretion of neurotrophic factors by neighboring endothelial cells (90–92). Angiogenic factors like VEGF-A and VEGF-C were increased in response to MSC-EVs suggesting a neuroprotective phenotype of the treated microglia. This interplay of different cytokines and chemokines are largely dependent on the disease context and may influence the outcome of the immune response. Balancing effective response and minimal damage to surrounding cells are complex functions of the immune system and require deeper investigation.

In addition to secretion of neuroinflammation-relevant proteins, we assessed the intracellular proteome of microglia and identified gene sets that were differentially enriched in response to MSC-EV treatment; thus, further demonstrating the significance of their morphological response. Some of the identified gene sets enriched in response to MSC-EVs were associated with amino acid metabolism (e.g. phenylalanine), inflammation (e.g. IL-4 and IL-13 signaling) and morphology (e.g. semaphorin and RHO/Rac signaling). Impaired phenylalanine metabolism was observed in our cytokine-stimulated microglia controls and increased phenylalanine concentration has been shown to activate microglia and alter their cytoskeleton due to an increase in their Iba-1 expression and size (93). MSC-EV treatment shows enrichment for phenylalanine metabolism in stimulated cells, reflecting a potentially desired therapeutic mechanism for addressing aberrant microglia activation in disease (94). Extracellular signaling proteins, such as semaphorins, are associated with changes in cell migration and motility for different cell-types in the nervous system (95) and proteins involved in this gene set were enriched in response to MSC-EV treatment. These results agree well with our observation of secreted proteins involved in regulation of cell motility and adhesion based on our STRING analysis. High enrichment score was observed for the IL-4 and IL-13 signaling gene set in MSC-EV treated groups. Reduced expression of disease associated microglia markers (e.g. TREM2, ApoE) have been reported following IL-4 and IL-13 treatment. In the same study, IL-4 also induced a metabolic shift in microglia towards oxidative phosphorylation (OXPHOS) (96). While commonly associated with anti-inflammatory response, these cytokines may augment neuroinflammation to promote influx of molecules that aid in repair and recovery (97). Increased levels of IL-4 along with CXCL8 in early AD patients was associated with a higher permeability of the BBB (98). Rho GTPase and Rac1 were among other pathways that were downregulated with stimulation and upregulated with MSC-EV treatment. These molecular switches are responsible for various cellular functions (e.g. proliferation, polarization) as well as morphology (e.g. reorganization of actin cytoskeleton, migration) (99). In microglia, RhoA balance is crucial to prevent neurodegeneration while simultaneously promoting microglial function (99–101). Additionally, microglia with reduced RhoA have modified actin stress fibers that was shown to induce an inflammatory phenotype (102).

Microglia lipid metabolism is dysregulated in different neurodegenerative diseases leading to lipid accumulation in cells (73). We observed significantly increased lipid content post stimulation which reflects the increased cell size (67) quantified using our morphological profiling approach. While PCA revealed separation of the EV treated group from +CTL group based on raw lipidomic feature data (**Figure S4)**, no significant changes in annotated lipids were observed in response to MSC-EV treatment (**Figure 5**). This could be explained by our relatively short (24 hour) experimental timeline and what is known about lipid metabolism dynamics. MSC-EV treatment effectively prevented the cytokine-induced morphological response of microglia, but 24 hours may not be sufficient for metabolism of accumulated lipids from the cell. Previous studies using immortalized mouse microglia cell-line BV2s (relevant to our work), as well as liver tissue, range from 18 – 72 hours to assess lipid degradation (66,103,104). Moreover, lipid metabolism is a multi-stage process and is dependent on lipid droplet size as observed in liver cells (103,105). It is possible that the regulation of microglia lipid content occurs over a longer period of time and thus temporal monitoring of lipidome is worth further investigation.

It is also important to consider MSC-EVs possess a lipid bilayer and (100) internalization of MSC-EVs through endocytosis, phagocytosis or attachment to microglia surface could be potentially contributing to the observed results (106). In our own published work, we have shown that MSC-EVs contain many of the same lipid classes (e.g. PCs, PEs, sphingomyelins, and PEPs) that were detected in microglia in this study (107). Vanherle et al. have effectively shown that cholesterol from macrophage-derived EVs support the remyelination and other regenerative functions of brain macrophages (108). Our results could be interpreted as differential lipid density/organization (i.e. lipids/area or lipids/volume) as we observed the same increased lipid accumulation in MSC-EV treated microglia as +CTL microglia, but that increase was not associated with increased cell size. This can be reconciled based on knowledge that lipids (e.g. cholesterol, sphingolipids, ceramides) are instrumental in regulation of membrane mechanics/fluidity and organization. Cell plasma membrane consists of differently saturated lipid classes which regulate the intercellular transport of signaling molecules as well as cellular morphology through actin re-organization (109). Like our observation, studies have shown microglia stimulation using other inflammatory signals (e.g. lipopolysaccharide, LPS) alters their lipid content (110) and leads to increased lipid raft formation (111) which induce morphological changes (112,113). Studies have also shown that microglia cells with lipid accumulation activate different pathways (e.g. LXR, PPAR) thus influencing cellular function(114). Further insight could be obtained through advanced single-cell based lipidomic approaches (e.g. MALDI based imaging (110,115)) as limitations exist with population wide approaches when considering heterogeneous cell populations (like microglia). Additionally, it will be critical to parse out the observed changes in lipid content and whether they are associated with the microglia or from lipids derived from the MSC-EVs, which could be achieved using isotopic labeling (115–117).

Although no significant overall changes in lipids were observed upon MSC-EV treatment, our imaging approach enabled assessment of another important regulator of microglia metabolism: mitochondria. An abundance of literature demonstrates the role and response of microglial mitochondria to neuroinflammation; for example, primary microglia stimulated with LPS show a significant increase in mitochondria fragmentation (118,119). This change in mitochondrial morphology is accompanied by metabolic reprogramming from OXPHOS to glycolysis (119,120). We observed that mitochondrial morphology in microglia is altered following cytokine stimulation and mitigated by MSC-EV treatment in a similar manner as overall cellular morphology. This further underscores the ability of morphological profiling to capture another important change in microglia phenotype relevant to their role in disease. Microglia mitochondrial dysregulation occurring due to either reduced expression of TREM2 (121), increased P2X_7_R expression (122), reduced TSPO-hexokinase2 activity (123) or impaired mitophagy (124) have been extensively reviewed in the context of Alzheimer’s disease (125–127). Further refinement of this assay could help investigate these mechanisms by additional mitochondrial membrane dyes (JC-1 (128), TMRM (129,130)) and/or more advanced high resolution microscopy approaches like confocal microscopy, scanning electron microscopy and STED super resolution microscopy that can profile mitochondria on a single-organelle level (albeit at significantly lower throughput) (124–129).

## 5. Conclusion

We demonstrated that significant overall changes in microglia morphology can be confidently interpreted to implicate the ability of MSC-EVs to modulate a multitude of disease-relevant microglia behaviors through comprehensive profiling of microglia mitochondrial morphology, lipidomics, secretion, and proteomics. The response of microglia to MSC-EVs suggest that MSC-EVs can exert a profound impact on the overall brain immune environment with microglia serving as a critical mediator of their therapeutic function. This is captured by changes in cellular and secreted proteins involved in pathways associated with inflammation, regulation of vasculature and other immune cells (e.g. macrophage migration, T-cell chemotaxis), which in turn may be responsible for regulation of the BBB. To explore these interactions further, we need to employ more complex assays involving neurons, endothelial cells and other glial cells using co-culture models (131–133) and organ-on-chip models with MSC-EV treatment (134–136).

This study established microglia morphology as an effective readout of MSC-EV bioactivity from a single donor and manufacturing protocol, and thus lays the foundation for using this assay to assess other key aspects of MSC manufacturing (48–50). Moving forward, this assay can be employed to screen different MSC cell-lines, manufacturing conditions (e.g. priming, media-type, effects of scaling) and EV long-term storage conditions (e.g. temperature, buffer). Additionally, this approach is amenable to high throughput screening and could thus be paired with compound libraries and genetic screens to elucidate MSC-EV mechanisms of action. Finally, this approach is generalizable and could be adapted to other MSC-EV target cells (besides microglia) that exhibit morphological-dependent behavior, as well as using this assay as a phenotypic readout of other therapeutics e.g. small molecules and other (non-MSC) cell-derived therapies.

## 6. Declarations

### Ethics Approval and Consent to Participate

Not Applicable.

### Consent for Publication

Not applicable.

### Availability of Data and Materials

The datasets used and/or analyzed during the current study are available from the corresponding author on reasonable request.

## Author’s Contributions

KRD: Conceptualization, Methodology, Software, Formal Analysis, Investigation, Writing – Original Draft, Review & Editing, Visualization AML: Methodology, Software, Investigation; MGM: Methodology, Investigation; KC: Methodology, Software; SB: Methodology, Software, Formal Analysis, Visualization, Writing – Review & Editing, JC: Methodology, Formal Analysis, Investigation, Writing – Review & Editing; HH: Methodology, Formal Analysis, Investigation, Writing – Review & Editing; KHM: Conceptualization, Formal Analysis, Resources, Writing – Review & Editing, Supervision, Project Administration Funding Acquisition; LBW: Conceptualization, Formal Analysis, Resources, Writing – Review & Editing, Supervision, Project Administration Funding Acquisition; RAM: Conceptualization, Methodology, Formal Analysis, Resources, Writing – Original Draft, Review & Editing, Visualization, Supervision, Project Administration, Funding Acquisition.

## Acknowledgments

Drs. Ty Maughon and Seth Andrews assisted with MSC expansion and cryopreservation for this work. Drs. Min-Ho Kim and Eric Dyne provided guidance for us to develop a microglia culture protocol. Thomas Spoerer, Courtney Campagna, Zhuofei Hou, Ravi Jyani, and Akash Ramachandran provided guidance in terms of experimental design, troubleshooting, and establishing the microglia morphology data pipeline. We would like to thank Dr. Rakesh Singh for processing the microglia samples for proteomic analysis, running the samples with mass spectrometry, and helping with interpretation and analysis of the annotated dataset. The authors thank Drs. Annie Bowles-Welch and Zhuoming Liu for their review of the manuscript.

