## Supplementary material for "Microglia Morphological Response to Mesenchymal Stromal Cell Extracellular Vesicles Demonstrates EV Therapeutic Potential for Modulating Neuroinflammation": supp figures

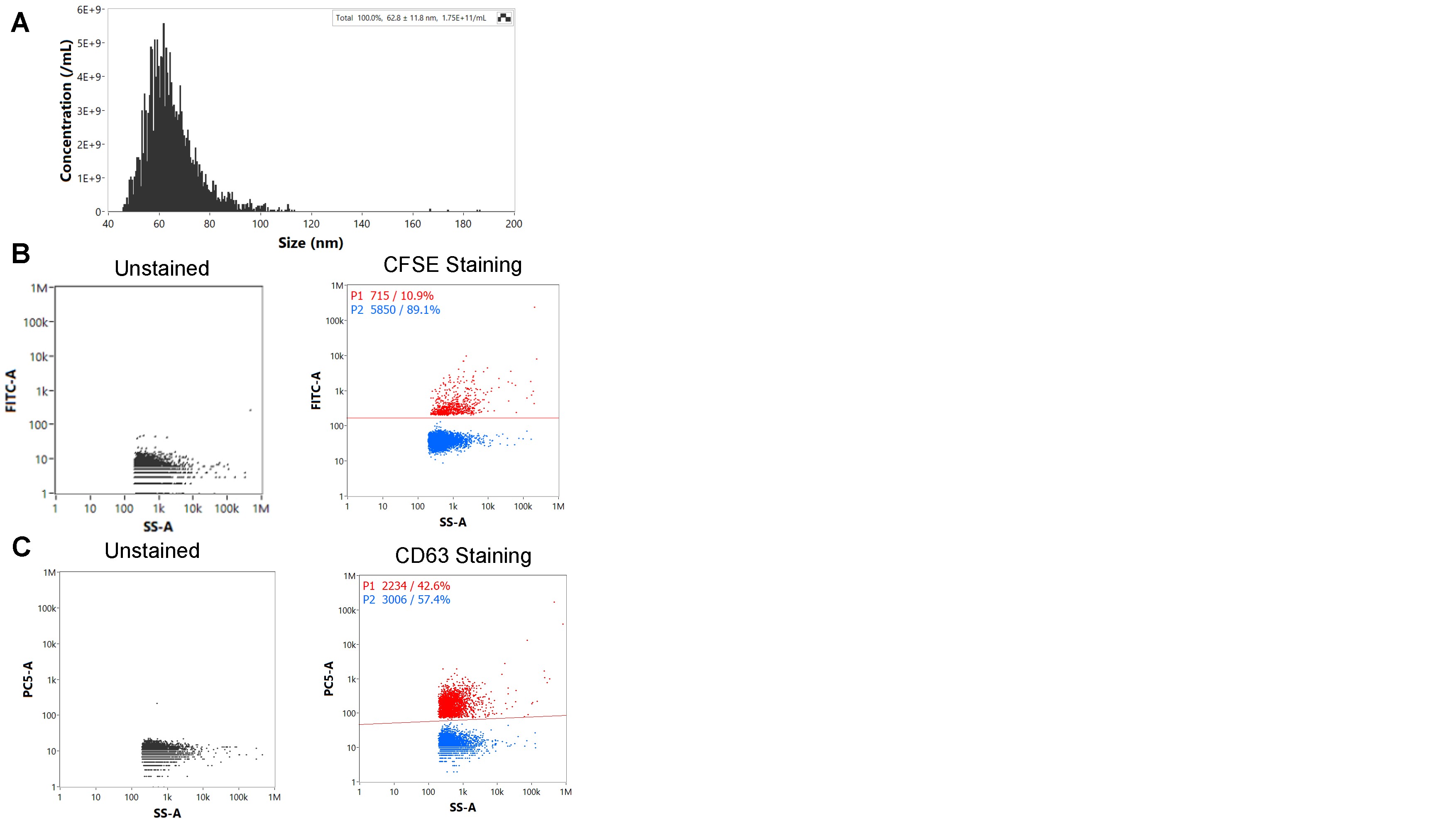

**Figure S1: MSC-EV surface marker staining  (A)** Concentration (particles/mL) and Size distribution (nm) of isolated EVs. Gating Strategy for single stains namely CFSE **(B)** and CD63 **(C)** conducted independently**.**

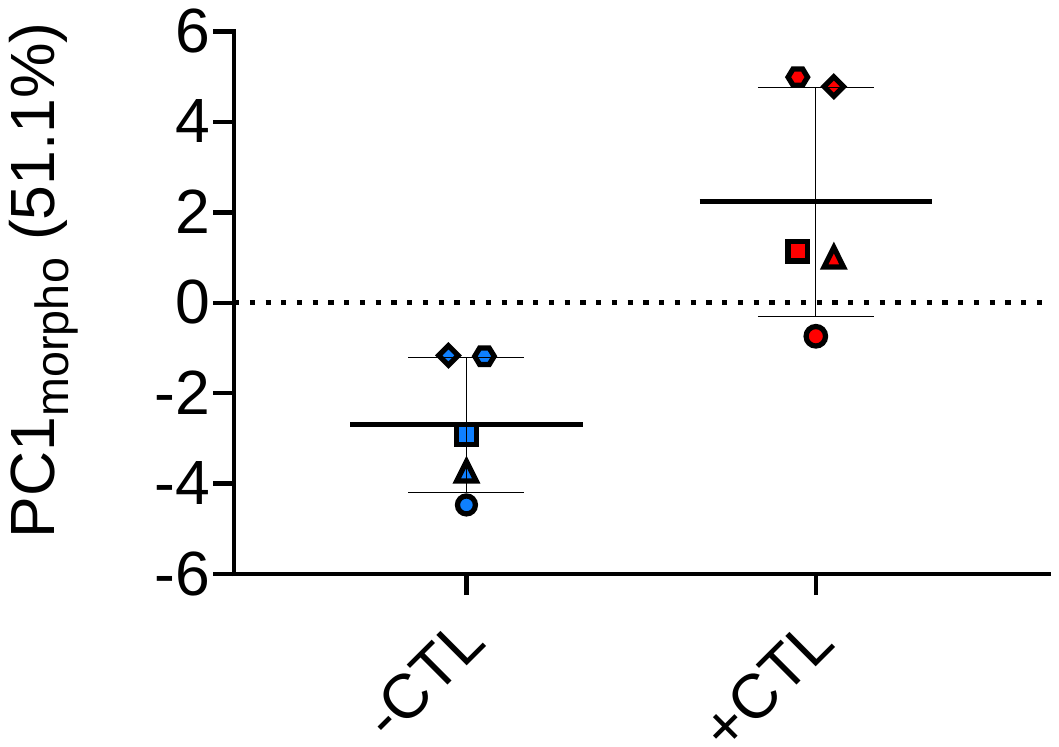
**Figure S2: Microglia morphology changes upon activation with cytokines across multiple experiments consistently.** Principal component 1 (PC1_morpho_) calculated using 21 features represents 51.1% of the variance between –CTL and +CTL. Each point is a mean 6 wells containing ~700-1000 cells/well with mean and standard deviation plotted for n=5 experiments per group. *p<0.01 vs -CTL for all groups conducted using paired t-test (two-tailed). Each experiment is represented by a different symbol.

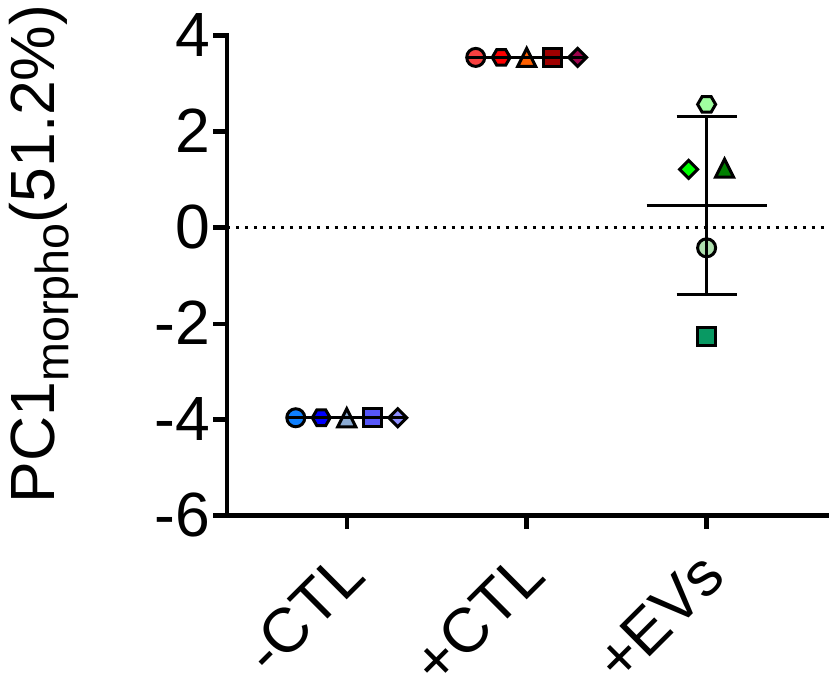
**Figure S3: Stimulated microglia morphology changes upon treatment with MSC-EVs from multiple batches.** Normalized principal component 1(PC1_morpho_) calculated using 21 min-max normalized features represents 51.2% of the variance. Each point is a mean of 6 wells calculated using median of ~700-1000 cells/well for n=5 experiments. *p<0.05 vs +CTL for all groups conducted using RM one-way ANOVA with multiple corrections using Dunnett multiple comparison testing.

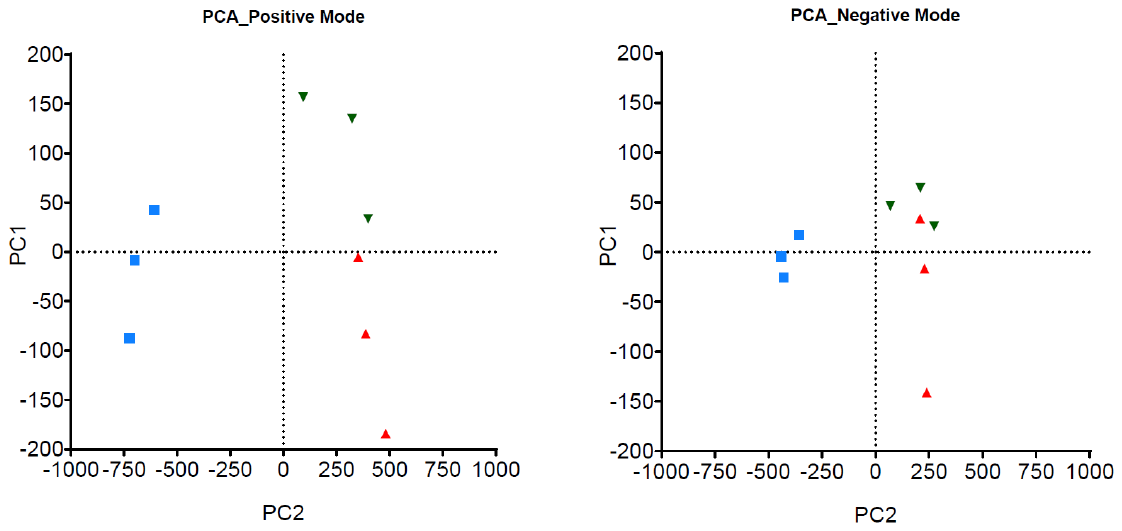
**Figure S4: PCA Plot of unannotated lipids.** Positive and Negative mode PCA plots of all lipids identified across the experimental groups –CTL, +CTL and +EVs with n=3 technical replicates.
